# Transcriptomic analysis of developing seeds in a wheat mutant RSD32 with reduced seed dormancy

**DOI:** 10.1101/846782

**Authors:** Kazuhide Rikiishi, Manabu Sugimoto, Masahiko Maekawa

**Affiliations:** Institute of Plant Science and Resources, Okayama University, Kurashiki, Okayama, Japan

## Abstract

Seed dormancy, a major factor regulating pre-harvest sprouting, can severely hinder wheat cultivation. Abscisic acid biosynthesis and sensitivity play important roles in the regulation of seed dormancy. Reduced Seed Dormancy 32 (RSD32), a wheat mutant with reduced seed dormancy, is derived from the pre-harvest sprouting tolerant cultivar, Norin61. RSD32 is regulated by a single recessive gene and mutant phenotype expressed in a seed-specific manner. Results of this study show that Norin61 has a low germination index (GI) of whole seeds at 50 days after pollination (DAP) and earlier developmental stages. In RSD32, higher GI of whole seeds was found on DAP40. Dormancy was released by DAP50. Gene expressions in embryos of Norin61 and RSD32 were compared using RNA-seq analysis at the different developmental stages of DAP20, DAP30, and DAP40. Numbers of up-regulated gene in RSD32 are similar in all developmental stages. However, down-regulated genes in RSD32 are more numerous on DAP20 and DAP30 than on DAP40. Homologous genes related to circadian clock regulation and Ca^2+^ signaling pathway, which have fundamental functions for plant growth and development, are involved in down-regulated genes in RSD32 on DAP20. For central components affecting the circadian clock, genes homologous to *CIRCADIAN CLOCK ASSOCIATED 1* (*CCA1*) and *LATE ELONGATED HYPOCOTYL* (*LHY*), which act as morning expressed genes, are expressed at lower levels in RSD32. However, higher expressions of *TIMING OF CAB EXPRESSION 1* (*TOC1*) and *PHYTOCLOCK 1* homologues, acting as evening expressed genes, are observed in RSD32. Homologues of Ca^2+^ signaling pathway related genes are found to be specifically expressed on DAP20 in Norin61. Lower expression is shown in RSD32. These results suggest that RSD32 mutation expresses on DAP20 and earlier seed developmental stages and suggest that circadian clock regulation and Ca^2+^ signaling pathway are involved in regulating wheat seed dormancy.

## Introduction

Wheat, similarly to rice and maize, is a major crop worldwide and is important for world food supplies. Pre-harvest sprouting, which is triggered by continuous rainfall during seed development, is seed germination on mother plants. This germination decreases seed quality and causes extensive economic damage to cultivation efforts. Seed dormancy is a major regulation factor affecting pre-harvest sprouting. Seed dormancy inhibits germination of seeds under favorable conditions (e.g. temperature and moisture). Nevertheless, it occurs after seed maturation has been completed [1]. Therefore, enhancing seed dormancy is an important breeding objective for avoiding pre-harvest sprouting damage.

Seed dormancy is induced and developed during seed maturation. Embryo development is completed at the early developmental stage (around 15 days after pollination: DAP15) in wheat [2]. Endosperms develop at the middle stage (DAP15–DAP30). The seed fresh weight reaches its maximum. Seeds reveal soft dough. At the late developmental stage (DAP30–DAP50), seed moisture contents decrease. Endosperms reach the hard dough stage. Seeds desiccate and change color from yellow to brown. Seed dormancy develops during seed desiccation in the late developmental stage.

A phytohormone, abscisic acid (ABA), plays an important role in the control of seed dormancy. In *Arabidopsis*, many mutants related to seed dormancy have been isolated. Earlier studies have indicated that ABA biosynthesis, catabolism, and sensitivity are involved in regulating seed dormancy [3–9]. *DELAY OF GERMINATION 1* (*DOG1*) has been identified as a quantitative trait locus (QTL) controlling the natural variation of seed dormancy in *Arabidopsis* [10]. In fact, *DOG1* interacts with ABA signaling pathway through type 2C protein phosphatases *ABA-HYPERSENSITIVE GERMINATION 1* (*AHG1*) and *AHG2* [11, 12]. Seed maturation regulators *LEAFY COTYLEDON 1* (*LEC1*), *LEC2*, *FUSCA3* (*FUS3*) and *ABA INSENSITIVE 3* (*ABI3*) are also involved in the regulation of seed dormancy [13–19]. These genes express at the early to late developmental stages of seed in *Arabidopsis*. In monocot species, *MOTHER OF FT AND TFL1* (*MFT*) and *MAP KINASE KINASE* in wheat [20, 21], *SDR4* in rice [22], and *ALANINE AMINOTRANSFERASE* (*AlaAT*)in barley [23] have been identified as QTLs regulating seed dormancy. Rikiishi et al. [24] reported that *TaABF1* related with ABA signaling pathway regulates the varietal variation of seed dormancy in wheat cultivars. Several genes regulating seed dormancy have been identified in wheat as well. These genes express at the late seed developmental stage. However, wheat genes homologous to *DOG1* and seed maturation regulators are expressed at the early to middle seed development stage [25]. The appropriate time for expression differs depending on the regulator gene. These results indicate that different regulatory systems function at each developmental stage for seed dormancy regulation.

Reports of the literature describe that ABA signaling is connected with and integrated with other signaling pathways. Calcium ion acts as a second messenger. In fact, calcium signals are involved in several stress responses to cold, drought, salinity and light in plants [26, 27]. The Ca^2+^ signaling pathway is initiated with the acceptance of Ca^2+^ signals by sensor proteins. Plant Ca^2+^ sensors belong to three families [29–30]. Calmodulins (CaMs) and CaM-like proteins (CMLs) are grouped in the same family. The second family is calcineurin B-like proteins (CBLs) that specifically activate CBL-interacting protein kinases (CIPKs). The third family is Ca^2+^-dependent protein kinases (CDPKs), which have a kinase domain and a Ca^2+^ sensor domain. Sensor proteins accepting Ca^2+^ signals are decoded to downstream responses. Because Ca^2+^ sensor proteins affect ABA sensitivity, Ca^2+^ signaling cooperatively regulates the response to stresses with the ABA signaling pathway [31–36]. Several reports have described the functions of circadian-clock-related genes on ABA sensitivity and dormancy release [37–41]. Circadian clock regulates the gene expressions and physiological responses corresponding to a daily cycle of light and darkness. In *Arabidopsis*, *LATE ELONGATED HYPOCOTYL* (*LHY*), *CIRCADIAN CLOCK ASSOCIATED 1* (*CCA1*), *TIMING OF CAB EXPRESSION1*/ *PSEUDO-RESPONSE REGULATOR* (*TOC1*/*PRR*), *EARLY FLOWERING* (*ELF*) and *LUX ARRHYTHMO* (*LUX*) are involved in the central cores of the circadian clock [38]. These genes encode transcription factors or proteins forming complexes with transcription factor and construct a complex system with feedback loop regulation. Circadian rhythm is oscillated by the interactions between the morning expressed *LHY* and *CCA1* and the evening expressed *TOC1*. Circadian clock components interact with various transcription factors such as *NIGHT LIGHT-INDUCIBLE AND CLOCK-REGULATED 1* (*LNK1*), *FAR-RED IMPAIRED RESPONSE 1* (*FAR1*), *PHYTOCHROME INTERACTING FACTORs* (*PIF*s), *REVEILLE*s (*RVE*s), and *CONSTANS-LIKE* [42–46]. Furthermore, circadian clock genes regulate cytosolic Ca^2+^ influx and the signaling pathways of ABA and Ca^2+^, suggesting that integrating regulatory network is necessary for circadian clock related fundamental processes in plant growth and development [47–50].

Rikiishi and Maekawa [51] produced a mutant with reduced seed dormancy from Norin61, a pre-harvest sprouting tolerant cultivar. Reduced Seed Dormancy 32 (RSD32) was found to be a seed-specific and single recessive mutation. Expressions of several transcription factors related with the regulation of seed dormancy were decreased in embryos of RSD32, suggesting that RSD32 is an important factor for the regulation of seed dormancy in wheat.

For the present study, gene expressions in embryos of Norin61 and RSD32 were investigated using RNA-seq analysis. Expression profiles were compared at three developmental stages: DAP20, DAP30, and DAP40. Results show that RSD32 mutation exhibits superior inhibitory effects on gene expression in embryos on DAP20 and DAP30. In embryos of RSD32, homologous genes of circadian clock and Ca^2+^ signaling pathway related genes were expressed differently from Norin61. RSD32 is a regulatory factor for wheat seed dormancy expressed at the middle developmental stage. The reduced seed dormancy in RSD32 might result from aberrant signals of circadian clock and Ca2+.

## Materials and Methods

### Plant materials and growth conditions

This study used a pre-harvest sprouting tolerant wheat cultivar, Norin61, and a mutant RSD32 with reduced seed dormancy selected from M_4_ population. They were derived from mutagenized Norin61 seeds using NaN_3_ treatment [51]. Seeds were sown in plastic trays for 4 weeks: 20 seedlings were transplanted to the field in each line with 20 cm between plants and 90 cm between rows. Plants were grown under a plastic roof to avoid rainfall. Spikes were tagged at anthesis. Seeds were harvested every 10 days from 10 days after pollination (DAP10) to DAP70 and were used in germination tests and RNA-seq analysis. To minimize variation, seeds were collected only from primary and secondary florets of the center spikelets.

### Germination test

Ten whole seeds were placed on filter paper in a Petri dish containing 6 ml of distilled water. Seeds were cut transversely into halves. Then ten half seeds with involved embryos were placed in a Petri dish containing 6 ml of distilled water with or without (±) 10 μM of ABA (Sigma Chemical Co.). The Petri dishes were then incubated in the dark at 20°C. All germination tests used three replications. Germinated seeds were counted daily for 14 days. A weighted germination index (GI) was calculated to give maximum weight to seeds that germinated first and to give less weight to those that germinated subsequently, as described by Walker-Simmons and Sesing [52]. GI values were converted into arcsine-transformed values and were used for statistical analyses.

### RNA isolation, library preparation and sequencing

Total RNA was isolated from three embryos on DAP20, DAP30, and DAP40 using a commercial kit (FastRNA Pro Green; Qbiogene Inc.). Isolated RNAs were purified using an RNA Clean-up Kit (TaKaRa Bio Inc., Tokyo, Japan). All kits were used according to the respective manufacturers’ protocols. The concentrations of total RNA samples were quantified using a spectrophotometer (Nano Drop ND-1000; Thermo Fisher Scientific Inc., Waltham, MA, USA). The quality of total RNA samples was also verified (Agilent 2100 Bioanalyzer; Agilent Technologies Inc., Santa Clara, CA, USA). The 18 RNA samples (2 lines × 3 stages × 3 biological replications) were sequenced. Library construction and sequencing for the Illumina HiSeq 2500 was provided as a custom service of Eurofins Genomics K.K. (Tokyo, Japan). After the polyA fraction (mRNA) was isolated from total RNA, it was fragmented. Then double-stranded (ds) cDNA was reverse-transcribed from the fragmented mRNA. The ds cDNA fragments were processed for adaptor ligation, size selection (for 200 bp inserts), and amplification to generate strand-specific cDNA libraries. Prepared libraries were subjected to paired-end 2 × 125 bp sequencing on the HiSeq 2500 platform with v4 chemistry.

### Bioinformatics analysis

We analyzed RNA-seq read data using RNA analysis tools in Galaxy/NAAC (https://galaxy.dna.affrc.go.jp/). Raw reads were obtained in Fastq format and were assessed for quality using FastQC. Terminal low-quality bases and adaptor sequences were trimmed off using Trimmomatic (Usadel Lab, Aachen, Germany). Clean reads were aligned against wheat survey sequence v3.0 obtained from International Wheat Genome Sequencing Consortium (IWGSC) using Tophat2 with default parameters. Cufflinks was used to assemble mapped reads. The resulting transcripts were used to quantify the expression of each gene in fragments per kilobase of transcript per million mapped reads (FPKM) unit. Cuffdiff was subsequently used to compile a list of differentially expressed genes (DEGs) with fold change ≥ 3 and *P*-value ≤0.01. BLASTX was used to align genes against the National Centre of Biotechnology Information (NCBI) database (https://blast.ncbi.nlm.nih.gov/Blast.cgi), the Rice Annotation Project Database (https://rapdb.dna.affrc.go.jp/tools/blast), Wheat Genetic Resources Database (https://shigen.nig.ac.jp/wheat/komugi/blast/blast.jsp) and Barley BioResources Database (http://earth.lab.nig.ac.jp/~dclust/cgi-bin/barley_pub/blast_search.html). The e-value cutoff was set at 1e-5. Gene names were assigned to each gene based on top Blastx hit with the highest score. RStudio (https://www.rstudio.com) was used for comparing of DEGs in three developmental stages.

## Results

### Seed dormancy and ABA sensitivity

Norin61 showed low germination indices (GIs) of whole seeds obtained on DAP50 and earlier stages. Strong seed dormancy was observed (Fig 1). GIs of half seeds, which were released dormancy, were reduced significantly with ABA incubation until DAP50. Norin61 showed sensitivity to ABA on seed germination. Seed dormancy and ABA sensitivity were lost after DAP60 in Norin61. However, RSD32 showed significantly higher GIs of whole seeds on DAP40 (50.0) and on DAP50 (90.5) than those in Norin61, although similar GIs of whole seeds were detected on DAP10–DAP30. RSD32 revealed the reduced seed dormancy phenotype at late developmental stages. GIs of half seeds were higher in RSD32 than those in Norin61 on DAP20 and DAP30, but no differences were observed on DAP40 and the later stages. Inhibitory effects of ABA on germination were reduced in RSD32 on DAP20, DAP30, and DAP50. These results indicate that RSD32 showed similar levels of seed dormancy to those of Norin61 on DAP20 and DAP30. However, seed dormancy was found to be reduced significantly on DAP40–DAP50 in RSD32. Reduced ABA sensitivity was also detected on DAP50 in RSD32.

**Fig 1.**
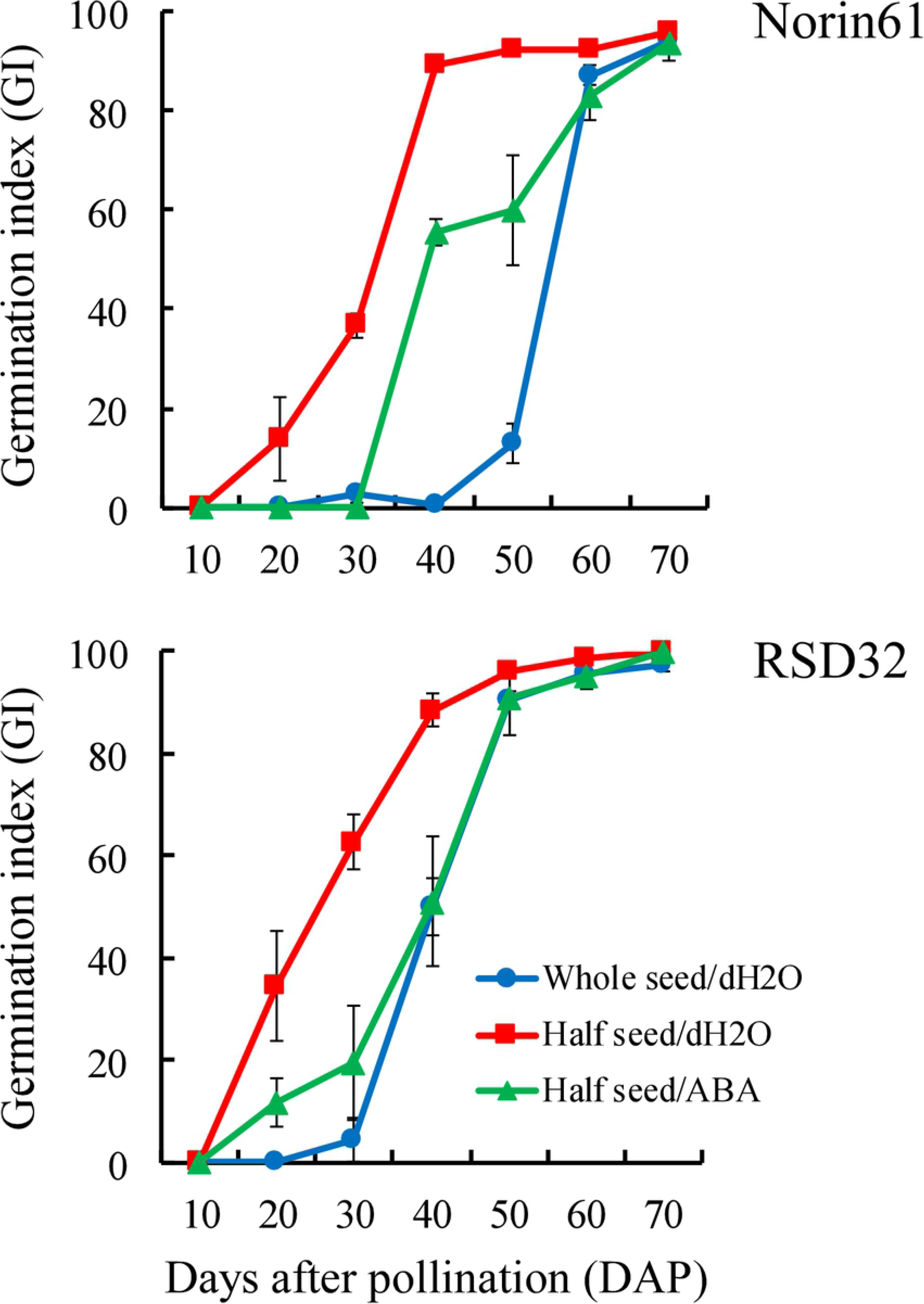
Germination index (GI) whole seeds in water and half seeds in water with and without 10 μM ABA in Norin61 and RSD32 at different developmental stages. Error bars represent SE.

### Differentially expressed genes (DEGs) during seed development

Reduction of seed dormancy was detected on DAP40 and DAP50 in RSD32. Gene expressions were compared at the middle to late developmental stages (DAP20, DAP30, DAP40) using RNA-seq analysis. Numbers of DEGs, which were down-regulated in RSD32, were 209, 228, and 49, respectively, on DAP20, DAP30, and DAP40 (Fig 2). Down-regulated genes in RSD32 were detected more on DAP20 and DAP30 than on DAP40. Numbers of DEGs, which were up-regulated in RSD32, were similar at all developmental stages. RSD32 mutation preferentially inhibited gene expression. Marked effects were observed at earlier developmental stages than on DAP40, when the seed dormancy reduction started.

**Fig 2.**
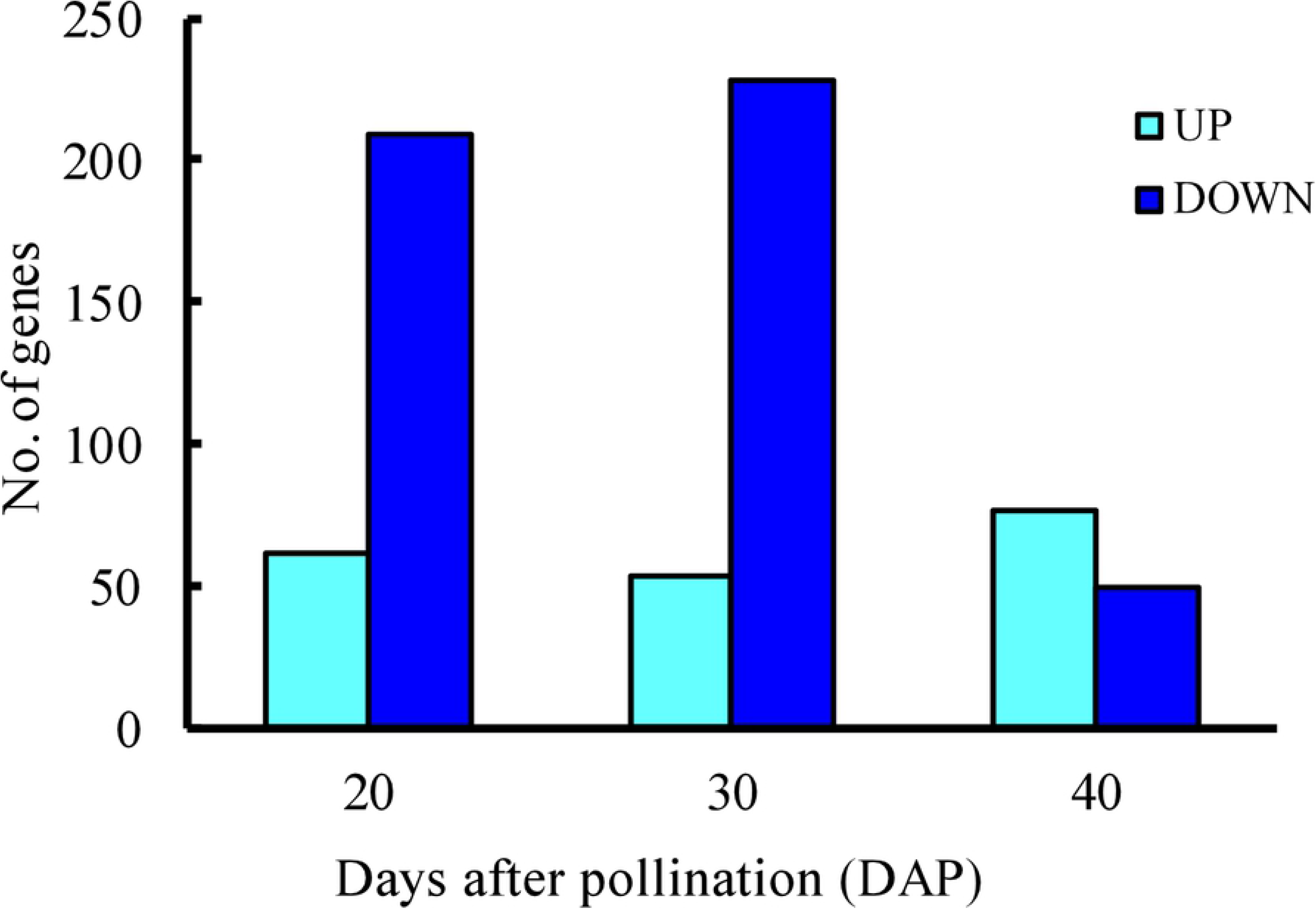
Numbers of differentially expressed genes (DEGs) between embryos of Norin61 and RSD32 at different developmental stages. UP, up-regulated genes; DOWN, down-regulated genes.

Comparison of down-regulated genes in RSD32 among developmental stages revealed that 146 and 164 DEGs showed specific expression on DAP20 and DAP30, respectively, and that 62 DEGs were down-regulated at both of DAP20 and DAP30 (Fig 3A). Most down-regulated genes on DAP40 showed stage-specific expression. No overlap with other developmental stages was observed. Up-regulated genes in RSD32 showed less overlap among developmental stages. Actually, most of the up-regulated genes were expressed specifically at each developmental stage (Fig 3B).

**Fig 3.**
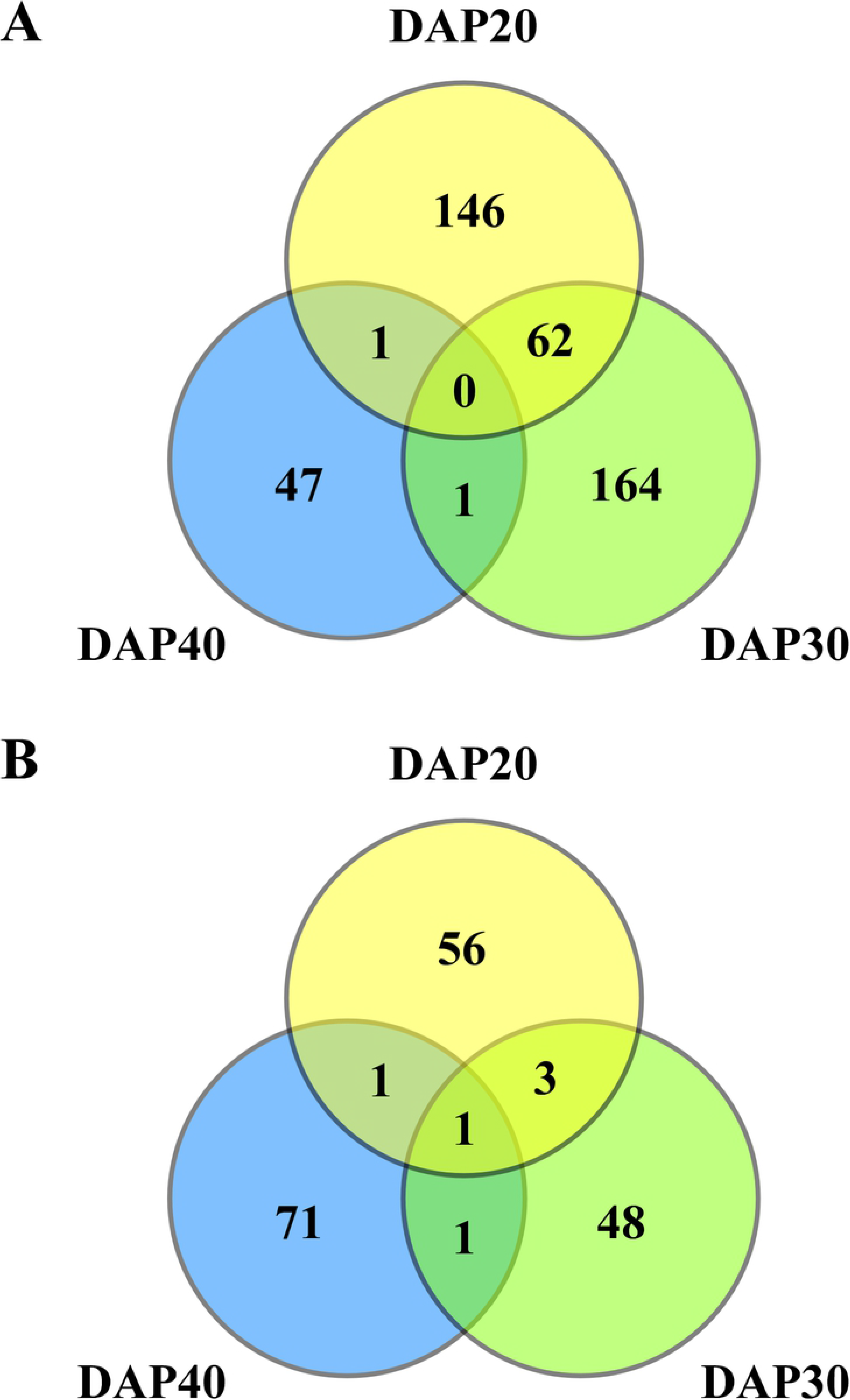
Venn diagram highlighting the number of differentially expressed genes in the three developmental stages. A, up-regulated genes; B, down-regulated genes.

At DAP20, down-regulated genes in RSD32 revealed several functions such as gene expression, protein metabolism, oxidation–reduction process, response to stimuli, circadian rhythm, and signal transduction (Tables 1 and 2). These genes were enriched homologous genes related to the calcium signaling pathway, such as *CALCIUM-BINDING PROTEIN*, *CALMODULIN-BINDING PROTEIN*, *EF-HAND Ca^2+^-BINDING PROTEIN*, *CALMODULIN-BINDING RECEPTOR-LIKE CYTOPLASMIC KINASE*, *CALCIUM-DEPENDENT PROTEIN KINASE*, and *CALCIUM-TRANSPORTING ATPASE*. These results indicate that unusual calcium signaling occurred in RSD32. Furthermore, down-regulated genes involved circadian-clock-related genes. Genes homologous to *CCA1*, *LHY*, *LNK1*, and *RVE6-LIKE* were expressed lower in RSD32 than those in Norin61. Circadian-clock-related genes were also identified as up-regulated genes in RSD32. Genes homologous to *FAR1-RELATED SEQUENCE 12-LIKE* and *CONSTANS-LIKE 9* were found to be expressed higher in RSD32 than in Norin61 on DAP20 (Table 3). A homologous gene to *TOC1* was also expressed 2.4 times higher in RSD32.

**Table 1.**
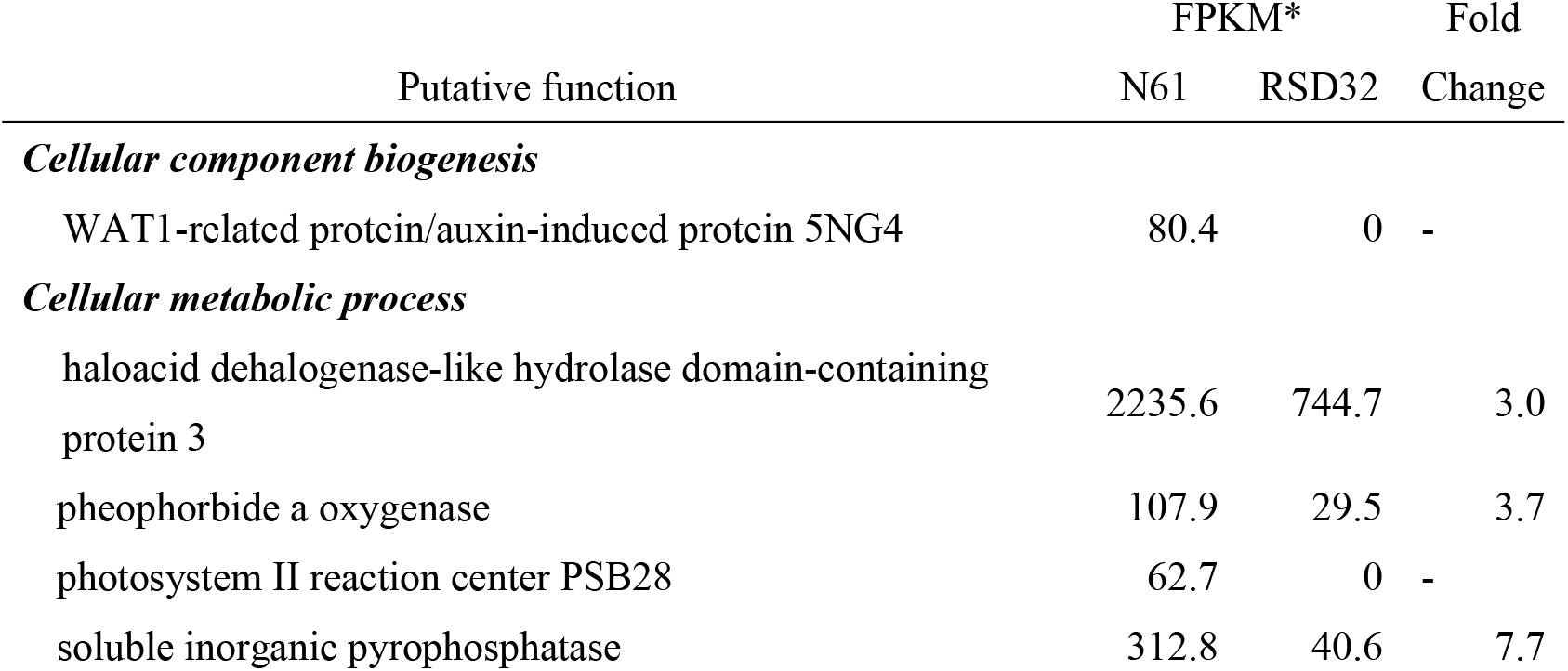
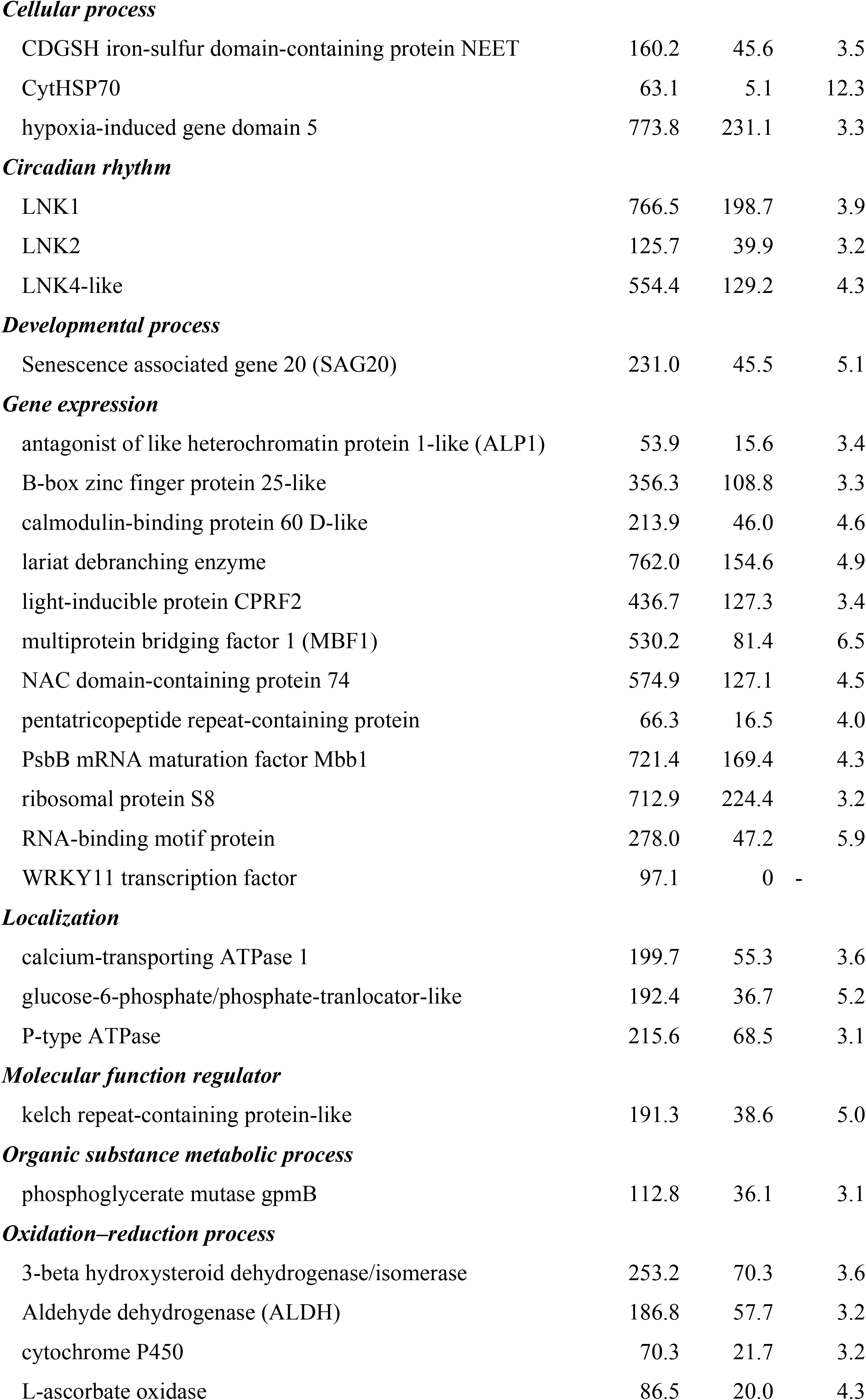
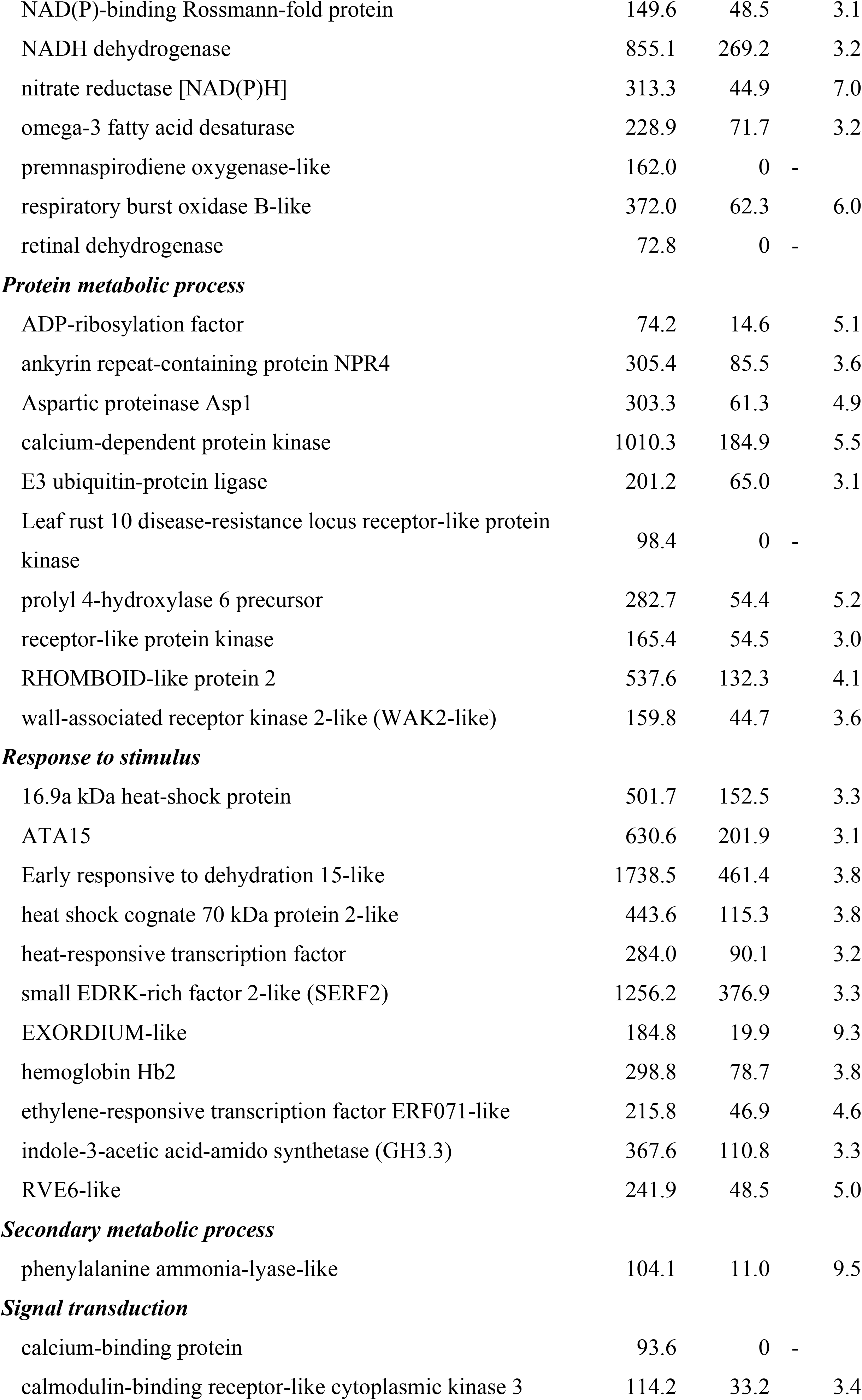
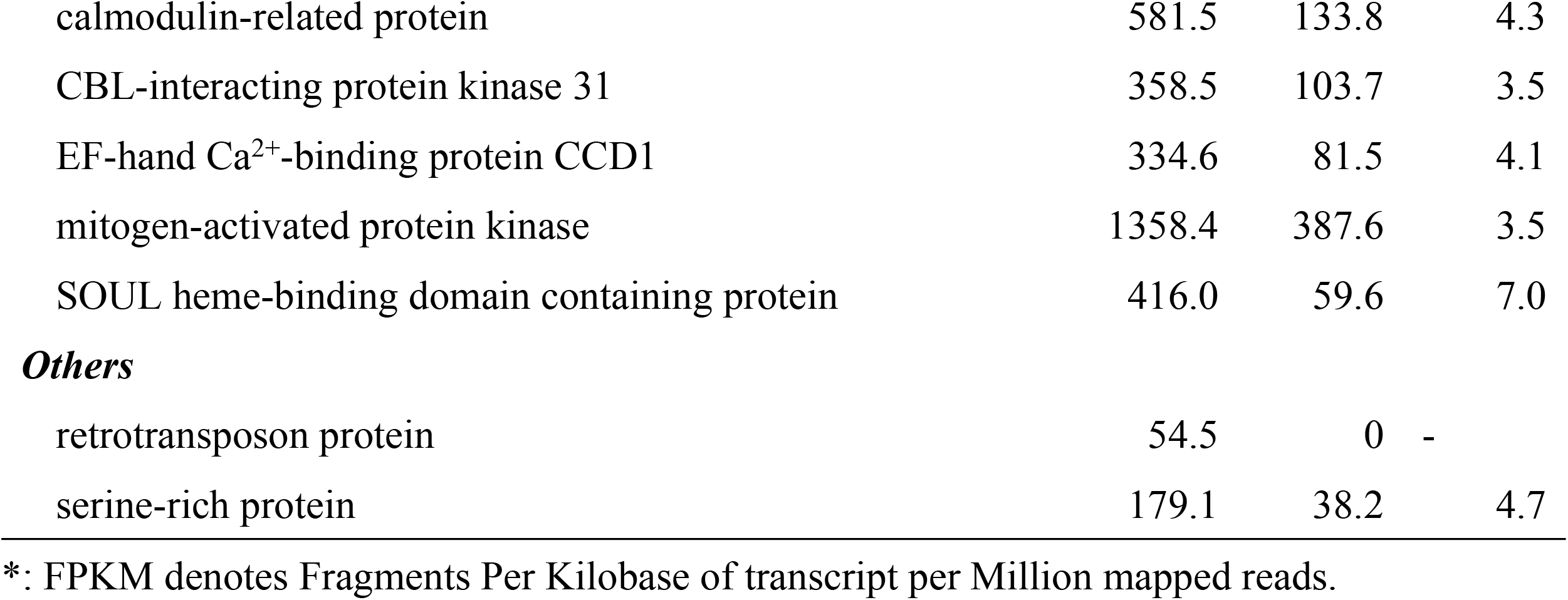
Down-regulated genes found on DAP20 in RSD32

| Putative function | FPKM* |  | Fold |
| --- | --- | --- | --- |
|  | N61 | RSD32 | Change |
| ***Cellular component biogenesis*** |  |  |  |
| WAT1-related protein/auxin-induced protein 5NG4 | 80.4 | 0 | - |
| ***Cellular metabolic process*** |  |  |  |
| haloacid dehalogenase-like hydrolase domain-containing protein 3 | 2235.6 | 744.7 | 3.0 |
| pheophorbide a oxygenase | 107.9 | 29.5 | 3.7 |
| photosystem II reaction center PSB28 | 62.7 | 0 | - |
| soluble inorganic pyrophosphatase | 312.8 | 40.6 | 7.7 |
| ***Cellular process*** |  |  |  |
| CDGSH iron-sulfur domain-containing protein NEET | 160.2 | 45.6 | 3.5 |
| CytHSP70 | 63.1 | 5.1 | 12.3 |
| hypoxia-induced gene domain 5 | 773.8 | 231.1 | 3.3 |
| ***Circadian rhythm*** |  |  |  |
| LNK1 | 766.5 | 198.7 | 3.9 |
| LNK2 | 125.7 | 39.9 | 3.2 |
| LNK4-like | 554.4 | 129.2 | 4.3 |
| ***Developmental process*** |  |  |  |
| Senescence associated gene 20 (SAG20) | 231.0 | 45.5 | 5.1 |
| ***Gene expression*** |  |  |  |
| antagonist of like heterochromatin protein 1-like (ALP1) | 53.9 | 15.6 | 3.4 |
| B-box zinc finger protein 25-like | 356.3 | 108.8 | 3.3 |
| calmodulin-binding protein 60 D-like | 213.9 | 46.0 | 4.6 |
| lariat debranching enzyme | 762.0 | 154.6 | 4.9 |
| light-inducible protein CPRF2 | 436.7 | 127.3 | 3.4 |
| multiprotein bridging factor 1 (MBF1) | 530.2 | 81.4 | 6.5 |
| NAC domain-containing protein 74 | 574.9 | 127.1 | 4.5 |
| pentatricopeptide repeat-containing protein | 66.3 | 16.5 | 4.0 |
| PsbB mRNA maturation factor Mbb1 | 721.4 | 169.4 | 4.3 |
| ribosomal protein S8 | 712.9 | 224.4 | 3.2 |
| RNA-binding motif protein | 278.0 | 47.2 | 5.9 |
| WRKY11 transcription factor | 97.1 | 0 | - |
| ***Localization*** |  |  |  |
| calcium-transporting ATPase 1 | 199.7 | 55.3 | 3.6 |
| glucose-6-phosphate/phosphate-tranlocator-like | 192.4 | 36.7 | 5.2 |
| P-type ATPase | 215.6 | 68.5 | 3.1 |
| ***Molecular function regulator*** |  |  |  |
| kelch repeat-containing protein-like | 191.3 | 38.6 | 5.0 |
| ***Organic substance metabolic process*** |  |  |  |
| phosphoglycerate mutase gpmB | 112.8 | 36.1 | 3.1 |
| ***Oxidation–reduction process*** |  |  |  |
| 3-beta hydroxysteroid dehydrogenase/isomerase | 253.2 | 70.3 | 3.6 |
| Aldehyde dehydrogenase (ALDH) | 186.8 | 57.7 | 3.2 |
| cytochrome P450 | 70.3 | 21.7 | 3.2 |
| L-ascorbate oxidase | 86.5 | 20.0 | 4.3 |
| NAD(P)-binding Rossmann-fold protein | 149.6 | 48.5 | 3.1 |
| NADH dehydrogenase | 855.1 | 269.2 | 3.2 |
| nitrate reductase [NAD(P)H] | 313.3 | 44.9 | 7.0 |
| omega-3 fatty acid desaturase | 228.9 | 71.7 | 3.2 |
| premnaspirodiene oxygenase-like | 162.0 | 0 | - |
| respiratory burst oxidase B-like | 372.0 | 62.3 | 6.0 |
| retinal dehydrogenase | 72.8 | 0 | - |
| ***Protein metabolic process*** |  |  |  |
| ADP-ribosylation factor | 74.2 | 14.6 | 5.1 |
| ankyrin repeat-containing protein NPR4 | 305.4 | 85.5 | 3.6 |
| Aspartic proteinase Asp1 | 303.3 | 61.3 | 4.9 |
| calcium-dependent protein kinase | 1010.3 | 184.9 | 5.5 |
| E3 ubiquitin-protein ligase | 201.2 | 65.0 | 3.1 |
| Leaf rust 10 disease-resistance locus receptor-like protein kinase | 98.4 | 0 | - |
| prolyl 4-hydroxylase 6 precursor | 282.7 | 54.4 | 5.2 |
| receptor-like protein kinase | 165.4 | 54.5 | 3.0 |
| RHOMBOID-like protein 2 | 537.6 | 132.3 | 4.1 |
| wall-associated receptor kinase 2-like (WAK2-like) | 159.8 | 44.7 | 3.6 |
| ***Response to stimulus*** |  |  |  |
| 16.9a kDa heat-shock protein | 501.7 | 152.5 | 3.3 |
| ATA15 | 630.6 | 201.9 | 3.1 |
| Early responsive to dehydration 15-like | 1738.5 | 461.4 | 3.8 |
| heat shock cognate 70 kDa protein 2-like | 443.6 | 115.3 | 3.8 |
| heat-responsive transcription factor | 284.0 | 90.1 | 3.2 |
| small EDRK-rich factor 2-like (SERF2) | 1256.2 | 376.9 | 3.3 |
| EXORDIUM-like | 184.8 | 19.9 | 9.3 |
| hemoglobin Hb2 | 298.8 | 78.7 | 3.8 |
| ethylene-responsive transcription factor ERF071-like | 215.8 | 46.9 | 4.6 |
| indole-3-acetic acid-amido synthetase (GH3.3) | 367.6 | 110.8 | 3.3 |
| RVE6-like | 241.9 | 48.5 | 5.0 |
| ***Secondary metabolic process*** |  |  |  |
| phenylalanine ammonia-lyase-like | 104.1 | 11.0 | 9.5 |
| ***Signal transduction*** |  |  |  |
| calcium-binding protein | 93.6 | 0 | - |
| calmodulin-binding receptor-like cytoplasmic kinase 3 | 114.2 | 33.2 | 3.4 |
| calmodulin-related protein | 581.5 | 133.8 | 4.3 |
| CBL-interacting protein kinase 31 | 358.5 | 103.7 | 3.5 |
| EF-hand Ca<sup>2+</sup>-binding protein CCD1 | 334.6 | 81.5 | 4.1 |
| mitogen-activated protein kinase | 1358.4 | 387.6 | 3.5 |
| SOUL heme-binding domain containing protein | 416.0 | 59.6 | 7.0 |
| **Others** |  |  |  |
| retrotransposon protein | 54.5 | 0 | - |
| serine-rich protein | 179.1 | 38.2 | 4.7 |

\*: FPKM denotes Fragments Per Kilobase of transcript per Million mapped reads.

**Table 2.**
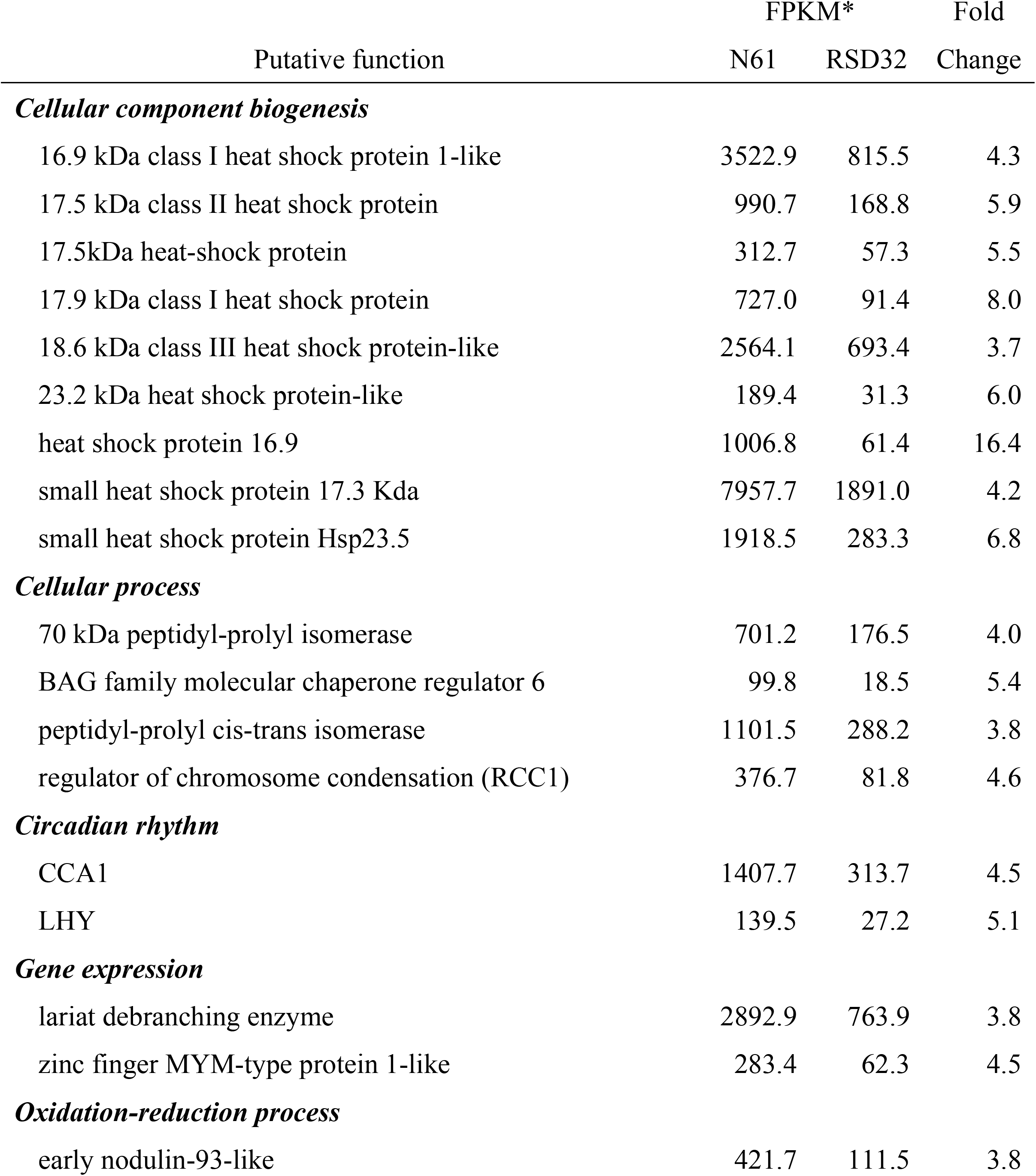
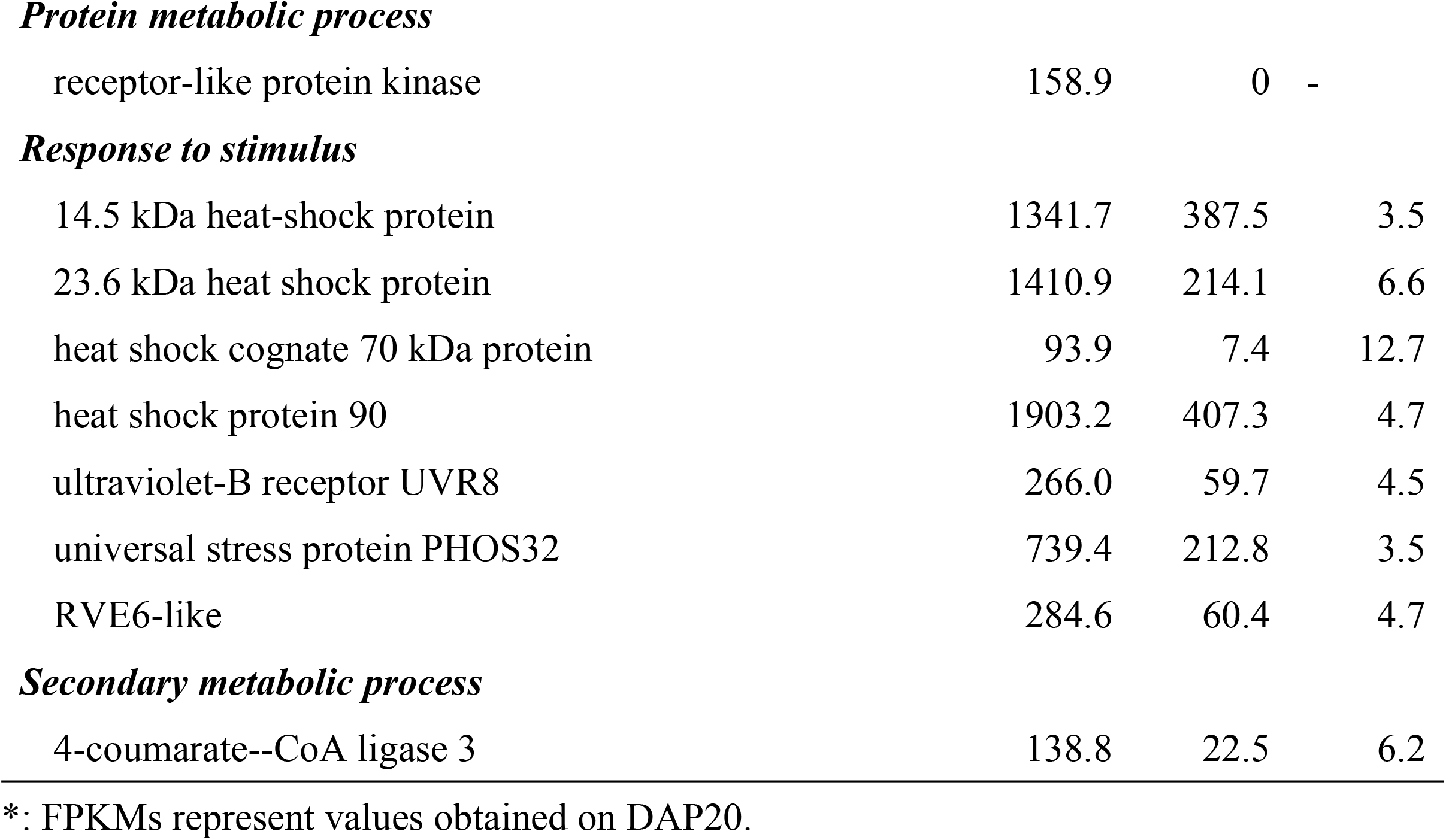
Down-regulated genes found on DAP20 and DAP30 in RSD32

| Putative function | FPKM* |  | Fold |
| --- | --- | --- | --- |
|  | N61 | RSD32 | Change |
| **Cellular component biogenesis** |  |  |  |
| 16.9 kDa class I heat shock protein 1-like | 3522.9 | 815.5 | 4.3 |
| 17.5 kDa class II heat shock protein | 990.7 | 168.8 | 5.9 |
| 17.5kDa heat-shock protein | 312.7 | 57.3 | 5.5 |
| 17.9 kDa class I heat shock protein | 727.0 | 91.4 | 8.0 |
| 18.6 kDa class III heat shock protein-like | 2564.1 | 693.4 | 3.7 |
| 23.2 kDa heat shock protein-like | 189.4 | 31.3 | 6.0 |
| heat shock protein 16.9 | 1006.8 | 61.4 | 16.4 |
| small heat shock protein 17.3 Kda | 7957.7 | 1891.0 | 4.2 |
| small heat shock protein Hsp23.5 | 1918.5 | 283.3 | 6.8 |
| **Cellular process** |  |  |  |
| 70 kDa peptidyl-prolyl isomerase | 701.2 | 176.5 | 4.0 |
| BAG family molecular chaperone regulator 6 | 99.8 | 18.5 | 5.4 |
| peptidyl-prolyl cis-trans isomerase | 1101.5 | 288.2 | 3.8 |
| regulator of chromosome condensation (RCC1) | 376.7 | 81.8 | 4.6 |
| **Circadian rhythm** |  |  |  |
| CCA1 | 1407.7 | 313.7 | 4.5 |
| LHY | 139.5 | 27.2 | 5.1 |
| **Gene expression** |  |  |  |
| lariat debranching enzyme | 2892.9 | 763.9 | 3.8 |
| zinc finger MYM-type protein 1-like | 283.4 | 62.3 | 4.5 |
| **Oxidation-reduction process** |  |  |  |
| early nodulin-93-like | 421.7 | 111.5 | 3.8 |
| **Protein metabolic process** |  |  |  |
| receptor-like protein kinase | 158.9 | 0 | - |
| **Response to stimulus** |  |  |  |
| 14.5 kDa heat-shock protein | 1341.7 | 387.5 | 3.5 |
| 23.6 kDa heat shock protein | 1410.9 | 214.1 | 6.6 |
| heat shock cognate 70 kDa protein | 93.9 | 7.4 | 12.7 |
| heat shock protein 90 | 1903.2 | 407.3 | 4.7 |
| ultraviolet-B receptor UVR8 | 266.0 | 59.7 | 4.5 |
| universal stress protein PHOS32 | 739.4 | 212.8 | 3.5 |
| RVE6-like | 284.6 | 60.4 | 4.7 |
| **Secondary metabolic process** |  |  |  |
| 4-coumarate--CoA ligase 3 | 138.8 | 22.5 | 6.2 |

\*: FPKMs represent values obtained on DAP20.

**Table 3.**
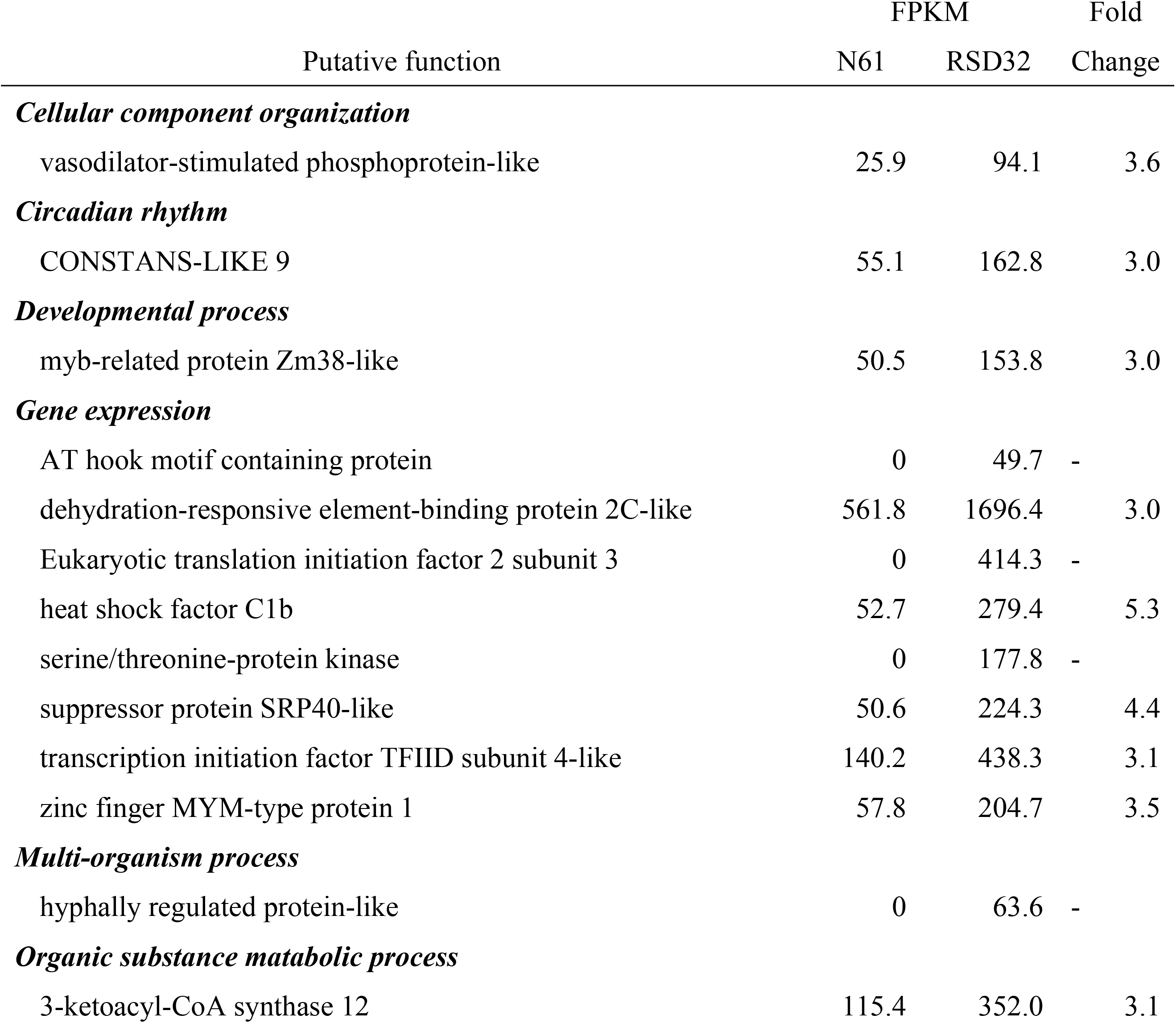
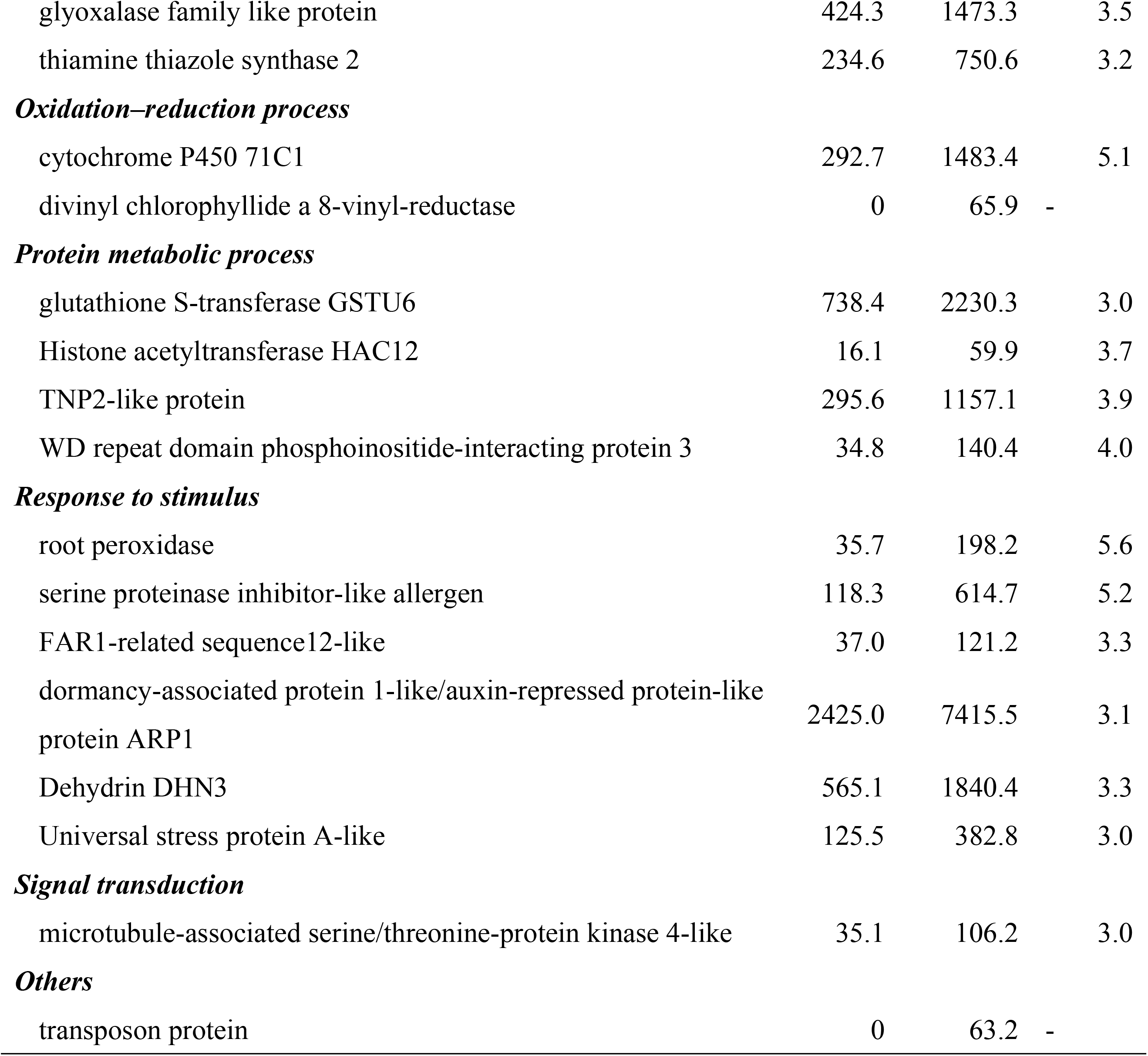
Up-regulated genes found on DAP20 in RSD32

| Putative function | FPKM |  | Fold |
| --- | --- | --- | --- |
|  | N61 | RSD32 | Change |
| **Cellular component organization** |  |  |  |
| vasodilator-stimulated phosphoprotein-like | 25.9 | 94.1 | 3.6 |
| **Circadian rhythm** |  |  |  |
| CONSTANS-LIKE 9 | 55.1 | 162.8 | 3.0 |
| **Developmental process** |  |  |  |
| myb-related protein Zm38-like | 50.5 | 153.8 | 3.0 |
| **Gene expression** |  |  |  |
| AT hook motif containing protein | 0 | 49.7 | - |
| dehydration-responsive element-binding protein 2C-like | 561.8 | 1696.4 | 3.0 |
| Eukaryotic translation initiation factor 2 subunit 3 | 0 | 414.3 | - |
| heat shock factor C1b | 52.7 | 279.4 | 5.3 |
| serine/threonine-protein kinase | 0 | 177.8 | - |
| suppressor protein SRP40-like | 50.6 | 224.3 | 4.4 |
| transcription initiation factor TFIID subunit 4-like | 140.2 | 438.3 | 3.1 |
| zinc finger MYM-type protein 1 | 57.8 | 204.7 | 3.5 |
| **Multi-organism process** |  |  |  |
| hyphally regulated protein-like | 0 | 63.6 | - |
| **Organic substance metabolic process** |  |  |  |
| 3-ketoacyl-CoA synthase 12 | 115.4 | 352.0 | 3.1 |
| glyoxalase family like protein | 424.3 | 1473.3 | 3.5 |
| thiamine thiazole synthase 2 | 234.6 | 750.6 | 3.2 |
| ***Oxidation–reduction process*** |  |  |  |
| cytochrome P450 71C1 | 292.7 | 1483.4 | 5.1 |
| divinyl chlorophyllide a 8-vinyl-reductase | 0 | 65.9 | - |
| ***Protein metabolic process*** |  |  |  |
| glutathione S-transferase GSTU6 | 738.4 | 2230.3 | 3.0 |
| Histone acetyltransferase HAC12 | 16.1 | 59.9 | 3.7 |
| TNP2-like protein | 295.6 | 1157.1 | 3.9 |
| WD repeat domain phosphoinositide-interacting protein 3 | 34.8 | 140.4 | 4.0 |
| ***Response to stimulus*** |  |  |  |
| root peroxidase | 35.7 | 198.2 | 5.6 |
| serine proteinase inhibitor-like allergen | 118.3 | 614.7 | 5.2 |
| FAR1-related sequence12-like | 37.0 | 121.2 | 3.3 |
| dormancy-associated protein 1-like/auxin-repressed protein-like protein ARP1 | 2425.0 | 7415.5 | 3.1 |
| Dehydrin DHN3 | 565.1 | 1840.4 | 3.3 |
| Universal stress protein A-like | 125.5 | 382.8 | 3.0 |
| ***Signal transduction*** |  |  |  |
| microtubule-associated serine/threonine-protein kinase 4-like | 35.1 | 106.2 | 3.0 |
| ***Others*** |  |  |  |
| transposon protein | 0 | 63.2 | - |

Down-regulated genes found on DAP30 were also enriched with several functions. Genes homologous to *PIF1-LIKE CCA1* and *LHY* were down-regulated in RSD32 (Table 2 and S1 Table). However, genes homologous to *PHYTOCLOCK1* and *LUX-B* showed higher expression in RSD32 on DAP30 (Table 4). Genes homologous to *CONSTANS-LIKE 9* and *FAR1-RELATED SEQUENCE 5-LIKE* were expressed, respectively, 2.7 times and 2.5 times higher in RSD32. Expressions of genes homologous to *CCA1* and *LHY* were inhibited in RSD32 at both DAP20 and DAP30. Down-regulated genes in RSD32 at both of DAP20 and DAP30 were enriched to *HEAT SHOCK PROTEIN*s (Table 2). Although some DEGs were identified as gene expression, oxidation–reduction process and protein metabolic process related genes on DAP40, circadian clock related genes were not found in DEGs on DAP40 (S2 and S3 Tables).

**Table 4.** Up-regulated genes found on DAP30 in RSD32

| Putative function | FPKM |  | Fold |
| --- | --- | --- | --- |
|  | N61 | RSD32 | Change |
| ***Cellular component biogenesis*** |  |  |  |
| WAT1-related protein | 0 | 34.7 | - |
| extracellular glycosidase CRH11-like | 0 | 391.6 | - |
| ***Cellular metabolic process*** |  |  |  |
| lipid phosphate phosphatase 3 | 18.8 | 80.0 | 4.3 |
| ***Circadian rhythm*** |  |  |  |
| LUX-B | 68.5 | 246.9 | 3.6 |
| PHYTOCLOCK1 | 161.1 | 536.4 | 3.3 |
| ***Developmental process*** |  |  |  |
| myosin-14-like | 0 | 110.3 | - |
| ***Gene expression*** |  |  |  |
| B3 domain-containing protein | 0 | 218.7 | - |
| trihelix transcription factor GTL1-like | 0 | 369.0 | - |
| zinc finger protein 410 | 0 | 8707.7 | - |
| ***Localization*** |  |  |  |
| sugar transporter ERD6-like 4 | 39.5 | 259.6 | 6.6 |
| ***Nucleic acid metabolic process*** |  |  |  |
| Superkiller viralicidic activity 2-like 2 | 259.5 | 868.1 | 3.3 |
| ***Organic substance metabolic process*** |  |  |  |
| plastid alpha-1,4-glucan phosphorylase | 10.8 | 48.9 | 4.5 |
| ***Oxidation–reduction process*** |  |  |  |
| premnaspirodiene oxygenase-like | 0 | 1048.3 | - |
| ***Protein metabolic process*** |  |  |  |
| deSI-like protein sdu1 | 40.9 | 185.7 | 4.5 |
| Lysine-specific demethylase 8 | 20.9 | 131.2 | 6.3 |
| RING-H2 finger protein | 0 | 3545.2 | - |
| Subtilisin-chymotrypsin inhibitor-2A | 262.7 | 913.9 | 3.5 |
| ***Response to stimulus*** |  |  |  |
| disease resistance protein RGA2-like | 282.2 | 872.3 | 3.1 |
| ***Signal transduction*** |  |  |  |
| Serine/threonine-protein kinase CTR1 | 91.6 | 352.7 | 3.9 |

### Temporal expression of homologous to circadian clock and Ca^2+^ signaling pathway related genes during seed development

Among DEGs, genes homologous to circadian clock and Ca^2+^ signal transduction related genes were abundant. In Norin61, genes homologous to *CCA1* and *LHY* showed the highest expression on DAP20. Their expressions were lower following seed development (Table 5). RSD32 showed lower expressions of *CCA1* and *LHY* than those of Norin61 on DAP20 or DAP30. However, genes homologous to *TOC1* expressed higher in RSD32 than those in Norin61 at all developmental stages. Homologous gene to *PHYTOCLOCK1* showed higher expression in RSD32 on DAP20 and DAP30, similarly to *TOC1*, but the difference was less apparent on DAP40. In other circadian clock related genes, genes homologous to *LUX-B* and *CONSTANS-LIKE* showed higher expressions in RSD32. Also, genes homologous to *LNK1*, *FAR1*, and *RVE6-LIKE* showed lower expression in RSD32. Consequently, RSD32 mutation was inferred to affect the expressions of circadian clock related genes in different manners. DEGs related to circadian clock regulation were divided into two groups based on mutant effects on the expression for inhibition or enhancement.

**Table 5.**
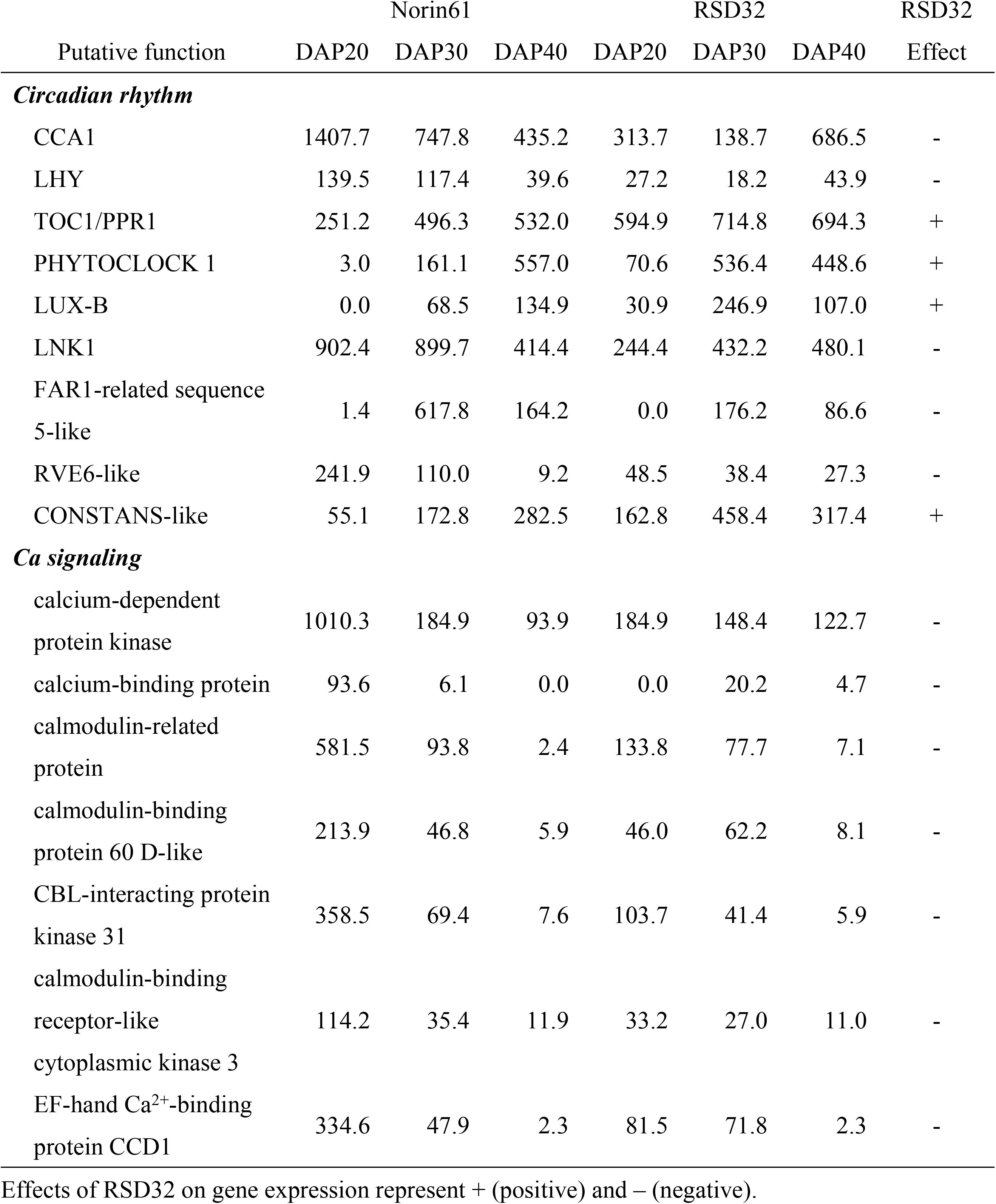
FPKMs of circadian rhythm and Ca signaling related genes at different developmental stages in Norin61 and RSD32

In Norin61, genes homologous to calcium signaling pathway related genes, *CALMODULIN-BINDING RECEPTOR LIKE CYTOPLASMIC KINASE 3*, *CALCIUM-BINDING PROTEIN*, *CALMODULIN-RELATED PROTEIN*, *CBL-INTERACTING PROTEIN KINASE 31*, *CALMODULIN-BINDING PROTEIN 60D-LIKE*, and *CALCIUM-DEPENDENT PROTEIN KINASE* were found to be specifically expressed on DAP20. Their expressions were found to be diminished on DAP30 and DAP40 (Table 4). These genes showed specific expressions at the middle developmental stage. Expressions of genes homologous to calcium signaling pathway related genes were found to be markedly inhibited on DAP20 in RSD32. All Ca signaling pathway related genes were down-regulated similarly.

## Discussion

Norin61 is a pre-harvest sprouting tolerant cultivar with strong seed dormancy. Although seed dormancy was maintained until DAP50, dormancy release was found on DAP60 and in later developmental stages. By contrast, RSD32 showed reduced seed dormancy on DAP40. Dormancy was found to be completely broken on DAP50. Levels of seed dormancy differed between Norin61 and RSD32 at the late developmental stages (DAP40 and DAP50). Both lines showed low germination ability in whole seeds. No differences were observed at middle developmental stages (DAP20 and DAP30). Because half seeds, which have been released from dormancy, showed poor germination, the germination ability was not fully developed at this stage. Transcriptome analysis of the gene expression in embryos of Norin61 and RSD32 at different developmental stages revealed that gene expressions were conspicuously different in both lines at the middle developmental stages, but not at the late developmental stages, which showed different levels of seed dormancy. These results suggest that RSD32 expresses at the middle developmental stage or earlier stage before seed dormancy development. Reduced seed dormancy in RSD32 is associated with genes expressed at the middle developmental stage.

Genes homologous to circadian clock regulation related genes are differentially expressed in embryos of Norin61 and RSD32 at the middle developmental stages. For the component of central oscillator, genes homologous to *CCA1* and *LHY* were down-regulated in RSD32. However, *TOC1* and *PHYTOCLOCK1* were up-regulated in RSD32. In *Arabidopsis*, *CCA1* and *LHY* are the morning expressed type; *TOC1* and *PHYTOCLOCK1* are the evening expressed type [38]. RSD32 affects the expressions of circadian clock regulation related genes depending on the circadian clock regulation function. Moreover, genes homologous to *LNK1*, *FAR1-RELATED SEQUENCE5-LIKE*, *RVE6-LIKE* and *CONSTANS-LIKE*, which interact with clock components, showed modified expressions in RSD32. In *Arabidopsis*, Penfield and Hall [37] report that circadian clock related genes were involved in dormancy release and that they affected the response to ABA and gibberellic acid (GA). Footitt et al. [40] reported that the balance between the evening and morning phases of the clock contributes to the interpretation of temperature signals, determining cycles of dormancy induction and relief in *Arabidopsis*. Aberrant function of the central oscillation affects ABA biosynthesis, signal transduction and several abiotic stress tolerances [38, 39, 41, 53–58]. The circadian clock might regulate several stress responses through ABA biosynthesis and the signal transduction pathway. Although the relation between circadian clock regulation and seed dormancy remains unknown in wheat, the reduction of seed dormancy in RSD32 might result from aberrant ABA signaling derived from irrelevant regulation of the circadian clock.

Genes homologous to calcium signaling pathway related genes were down-regulated in embryos of RSD32. Ca^2+^ sensor proteins, calmodulin-related protein, CDPK and CIPK, were identified as down-regulated genes in RSD32. In *Arabidopsis*, Ca^2+^ influx, and the expressions of *CALMODULIN-LIKE PROTEIN 39* (*CML39*), *CALCIUM DEPENDENT PROTEIN KINASE* (*CDPK*, *CPK*) and *CBL-INTERACTING PROTEIN KINASE* (*CIPK*) affect ABA signaling [31, 33, 35, 59–61]. In monocot species, *CIPK* and *CPK* also affect sensitivity to ABA [32, 34, 36]. Furthermore, Somyong et al. [62, 63] reported that the region-located wheat pre-harvest sprouting regulating QTL, QPhs.cnl-2B.1, involved several genes associated with Ca^2+^ signaling pathway, such as *CDPK*s and *CALMODULIN*/*Ca^2+^-DEPENDENT PROTEIN KINASE*. In this study, genes homologous to calcium signaling pathway related genes were found to be temporarily expressed in wheat embryos on DAP20: the middle developmental stage. Temporal induction of these genes was lost in RSD32. The seed dormancy induction might be disturbed by attenuated Ca signaling. Martí Ruiz et al. [50] reported that *CALMODULIN-LIKE 24* (*CML24*) regulates the expression of *TOC1* through Ca^2+^-dependent pathway in *Arabidopsis*. Calcium signaling might affect the expression of circadian clock related genes in wheat. Relations among seed dormancy, ABA signal transduction, circadian clock regulation and Ca^2+^ signaling remain unknown, but they should be investigated further especially in wheat. Few reports describe the functions of circadian clock and Ca^2+^ signaling on the regulation of wheat seed dormancy. RSD32 is a useful tool for investigating the complex network of these regulatory pathways in wheat.

Wheat embryo development is completed around 15 days after pollination (DAP15). Endosperms are developing continuously. In fact, the seed reaches maximum fresh weight at the middle developmental stage (DAP15–DAP30) [2]. Moisture contents of seeds are still high at this stage. At the late developmental stage (DAP30–DAP50), moisture contents of seeds are decreasing; seeds enter the dormant state. Physiological states differ between the seeds at the middle and late developmental stages. Although some DEGs were down-regulated in RSD32 at both DAP20 and DAP30, most DEGs identified on DAP40 are specifically expressed. No overlap with other developmental stages was observed. These results also indicate that seeds on DAP20 and DAP30 have similar physiological conditions, but they differ from those found on DAP40. Most studies investigating the regulation of seed dormancy have specifically examined the regulatory pathways of the late developmental stage. In monocot species, *MFT*, *MAP KINASE KINASE* and *AlaAT* have been identified as QTLs for regulating seed dormancy [20, 21, 23]. These genes are associated with maintenance and release of seed dormancy. They express at the late developmental stage. Because RSD32 expressed at the middle developmental stage, RSD32 might be an important gene for the regulation of seed dormancy, acting more upstream in the regulation pathway. In *Arabidopsis*, *DOG1* and seed maturation regulators function for the regulation of seed dormancy and express at early to middle developmental stages [10, 13–19]. Wheat genes homologous to *DOG1* and seed maturation regulators are also expressed at early to middle developmental stages [25]. Rikiishi et al. [51] reported decreased expression of *TaDOG1* in the embryos of RSD32. These results suggest that RSD32 acts upstream on *TaDOG1* function. Regulation factor expressed at the middle developmental stage might be associated with seed dormancy initiation. Although many studies have examined the development and maintenance of dormancy, the regulation mechanism for initiation and induction of dormancy remains unknown. Early events of seed dormancy regulation in wheat are elucidated by the identification of RSD32 function. Furthermore, understanding the relations among regulation systems expressed at different developmental stages is necessary to elucidate the overall network regulating seed dormancy.

## Supporting information

**S1 Table Down-regulated genes found on DAP30 in RSD32**

**S2 Table Down-regulated genes found on DAP40 in RSD32**

**S3 Table Up-regulated gens found on DAP40 in RSD32**

